# Time-restricted feeding from adulthood to old age improves biconditional associative learning in geriatric rats regardless of macronutrient composition

**DOI:** 10.1101/2020.02.06.937532

**Authors:** Abbi R. Hernandez, Quinten P. Federico, Sara N. Burke

## Abstract

Declining health and cognition are hallmarks of advanced age that reduce both the quality and length of the lifespan. While caloric restriction has been highlighted as a dietary intervention capable of improving the healthspan by restoring metabolic function in late life, time-restricted feeding and changes in dietary macronutrient composition may be more feasible alternatives with similar health outcomes. To investigate the potential of these two interventions, a pilot cohort of fully mature adult rats were placed on a time-restricted feeding regimen of a ketogenic or micronutrient and calorically matched control diet from 8 to 21 months of age. A third group of rats was permitted to eat standard chow *ad libitum*. At 22 months, all rats were then placed on time-restricted feeding and tested on a biconditional association task. While the data presented here are preliminary (small sample size of 3-4/diet group), additional animals are currently undergoing the feeding regimen, and will be added into the behavioral studies at a later date. For the current data, regardless of dietary composition, time-restricted-fed rats performed significantly better than *ad libitum*-fed rats. This observation could not be accounted for by differences in motivation, procedural or sensorimotor impairments, indicating that mid-life dietary interventions capable of preventing metabolic impairments may also serve to prevent age-related cognitive decline.

## Introduction

Two of the most prominent hallmarks of advancing age are declining peripheral health and impaired cognitive function, which are reciprocally linked (1). Caloric restriction, which has been shown to increase the lifespan in several species, has been posited to increase healthspan and cognitive function (2). The difficulty of maintaining long-term caloric restriction in humans, however, limits the translational potential of this lifestyle intervention for improving cognitive and physical function in older adults. Importantly, both time-restricted feeding (which is comparable to intermittent fasting) (3) and nutritional ketosis (4) mimic several aspects of caloric restriction and may confer health benefits to aged populations while imposing less severe dietary restrictions. A previous study reported that >3 months of nutritional ketosis, when initiated in aged rats (21 months old), resulted in improved cognitive function compared to rats that were on a standard carbohydrate-based control diet. Importantly, both diets had equivalent caloric and micronutrient content and were given once per day. Critically, the aged animals in this previous study had metabolic impairments, including hyperinsulinemia and excess visceral white adipose tissue (5). The possibility therefore exists that diet-induced cognitive benefits are directly related to the efficacy of different diets at reversing metabolic deficits that lead to impairments in brain function. Thus, it is important to determine if dietary interventions initiated in young adults, prior to the development of metabolic dysfunction, interacts with cognitive function in old age.

Age-related declines in cognitive performance can hinder quality of life in older adults. The goal of this study was to therefore investigate whether long-term time-restricted feeding initiated in young adulthood could improve cognitive outcomes in aged rats, and the extent to which this interacts with macronutrient composition. Two groups of rats were placed on a time-restricted feeding regimen beginning at 8 months of age. These rats were given ∼51 kcal of food once daily. All animals consumed the full ration of calories within 3 hours, resulting in ∼21 hours of fasting (5). Among the rats given time-restricted feeding, one group was fed a ketogenic diet, while the other group was fed a micronutrient and calorically equivalent control diet (6). A third group of rats was fed *ad libitum* until 21 months of age, at which time they were fed standard rodent chow once daily to encourage appetitively-motivated participation in cognitive testing. A previous study has reported that rats of the Fischer 344 × Brown Norway hybrid strain develop hyperinsulinemia and metabolic impairments when allowed unrestricted access to standard laboratory rodent chow from adulthood into old age (5).

In old age, all rats where tested on a biconditional associated task (BAT), which quantifies an animal’s ability to cognitively multitask by simultaneously alternating between two different arms of a maze while completing an object discrimination problem. Importantly, the correct choice of the target object updates based on the animal’s current location on the maze (Figure 1). Performance on this type of object-place paired associative learning has repeatedly been shown to decline with age in rats (7–10), and has greater sensitivity for detecting age-related impairments than the Morris watermaze test of spatial learning and memory (8). Critically, the BAT is more comparable to complex cognitive tasks that older human adults must complete for instrumental activities of daily living and therefore is a better behavioral metric for assessing the translational potential of novel interventions. Potential confounds due to differences in motivation, or procedural and sensorimotor impairments, were assessed with a simple object discrimination problem, in which performance is typically not impaired in aged rats (7).

**Figure 1:**
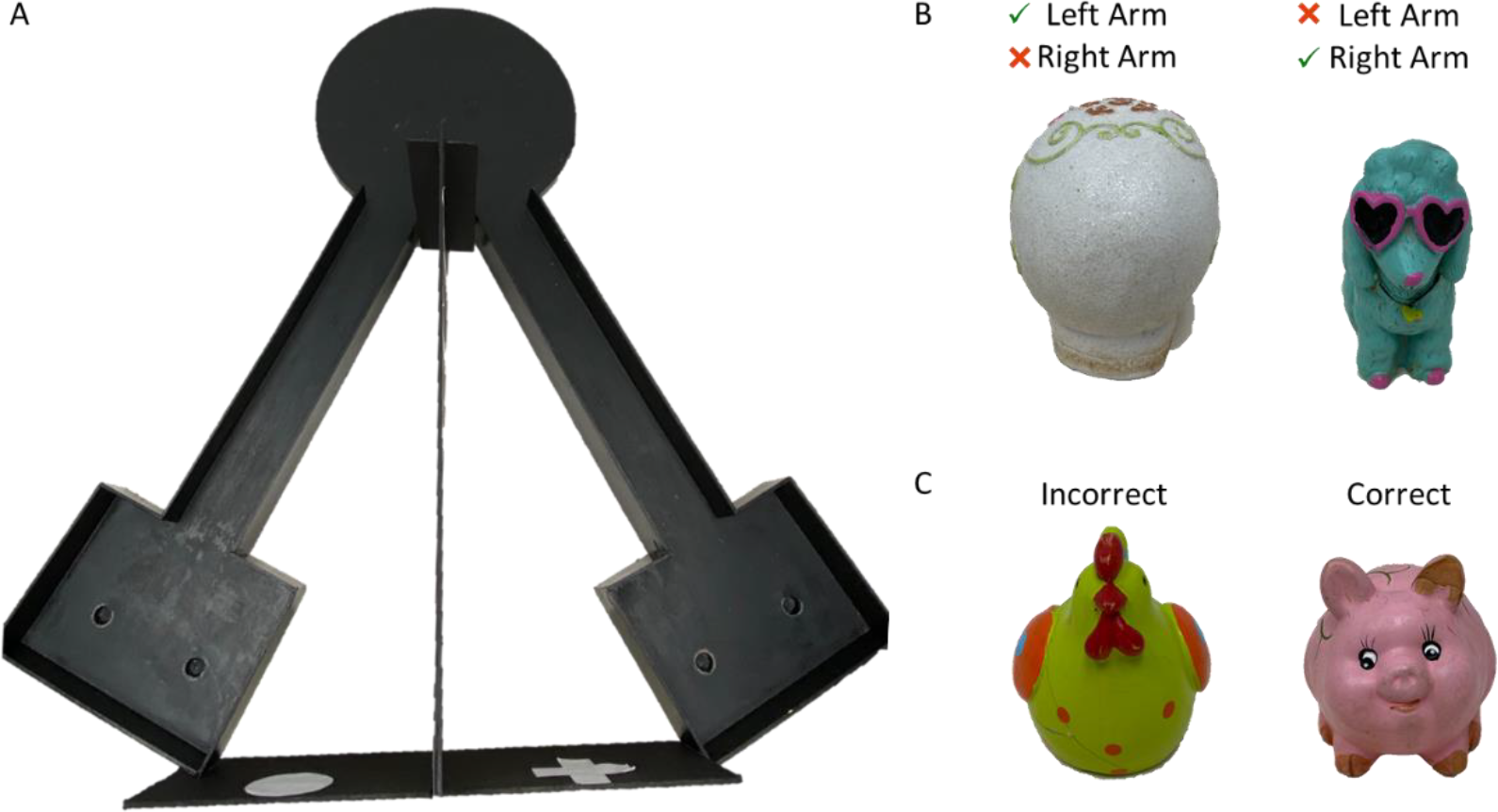
Behavioral testing apparatus and objects utilized for BAT and object discrimination tasks. (A) Birds-eye view of the testing apparatus used for alternations, BAT and object discrimination tasks. Note during the object discrimination, only one arm of the maze was used and rats were not required to alternate. Objects utilized during (B) BAT and (C) simple object discrimination testing.

## Methods

### Subjects and Dietary Interventions

11 aged (22 months) male Fisher 344 × Brown Norway F1 (FBN) Hybrid rats from the National Institute on Aging colony at Charles River were used in this study. All rats were housed individually and maintained on a 12-hour light/dark cycle with all behavioral testing occurring in the dark phase. Rats were divided into three groups: 1) fed *ad libitum* standard rodent chow until 21 months (n = 3), 2) fed 51 kCal of a standard diet once daily from months 8 to 21 (n = 4) and 3) fed 51 kCal of a ketogenic diet once daily from months 8 to 21 (n = 4). At 21 months of age, all rats were further restricted (approximately 25-30 kCal/day) to encourage participation in the appetitively-motivated BAT behavior. Water was provided to all rats *ad libitum* throughout the study.

The same ketogenic diet (KD) and micronutrient matched control diet (CD) were used as published previously (6,9,11). An additional group of rats were fed *ad libitum* with standard laboratory chow (Envigo, Teklad 2918). The KD was a high fat/low carbohydrate diet (Lab Supply; 5722, Fort Worth, Texas) mixed with MCT oil (Neobee 895, Stephan, Northfield, Illinois) with a macronutrient profile of 76% fat, 4% carbohydrates, and 20% protein. The micronutrient-matched CD (Lab Supply; 1810727, Fort Worth, Texas) had a macronutrient profile of 16% fat, 65% carbohydrates, and 19% protein. Nutritional ketosis was verified by testing peripheral levels of glucose and the ketone body β-hydroxybutyrate (BHB) 1 hour after feeding.

### Behavioral Testing

Rats were trained on the biconditional association task (BAT) as previously published (8,10,12). Briefly, rats were first trained to alternate between left and right arms of a V-shaped maze (see Figure 1) with a macadamia nut reward at the end of each arm. Alternation training continued until rats reached a criterion performance of ≥80% correct with completion of all 32 trials within 20 minutes. Failing to alternate was logged as an incorrect trial. Following alternation training, rats began testing on the BAT, in which a single object pair was placed over two different food wells in the choice platform at the end of both arms (Figure 1B, orb and blue poodle). One object covered a hidden food reward that the rat could retrieve for moving the correct object. Importantly, in the left arm the orb was the rewarded object while in the right arm the poodle was the rewarded object. Rats were allowed to eat the food reward (macadamia nut) if they correctly displaced the object contingent on the current location within the maze. Rats were given 32 trials per day in alternating arms with objects placed pseudorandomly on the left and right sides within a given arm. Rats were trained until a criterion performance of ≥ 80% correct for each object on 2 consecutive days. Following criterion performance on the BAT, all rats were tested on a simple object discrimination within a single arm of the maze. For this control task, object choice was not contingent upon location, and the same object was always rewarded. For both tasks involving objects, selecting the unrewarded object was logged as an incorrect trial. During this phase of testing, rats did not make any alternation errors.

## Results

Postprandial glucose (Figure 2A) and BHB (Figure 2B) measurements were utilized to generate a glucose ketone index (GKI) for each rat as reported previously (11). GKI values during behavioral testing indicate only rats fed the ketogenic diet were in nutritional ketosis (F_[2,8]_ = 49.62; p < 0.001; Figure 2C). There was no effect of feeding method (F_[1,9]_ = 0.78; p = 0.40), as rats on time-restricted feeding with the standard diet did not differ from *ad libitum*-fed rats (t_[5]_ = 1.45; p = 0.21), but had significantly higher GKI values than ketogenic-fed rats (t_[6]_ = 8.55; p < 0.001).

**Figure 2:**
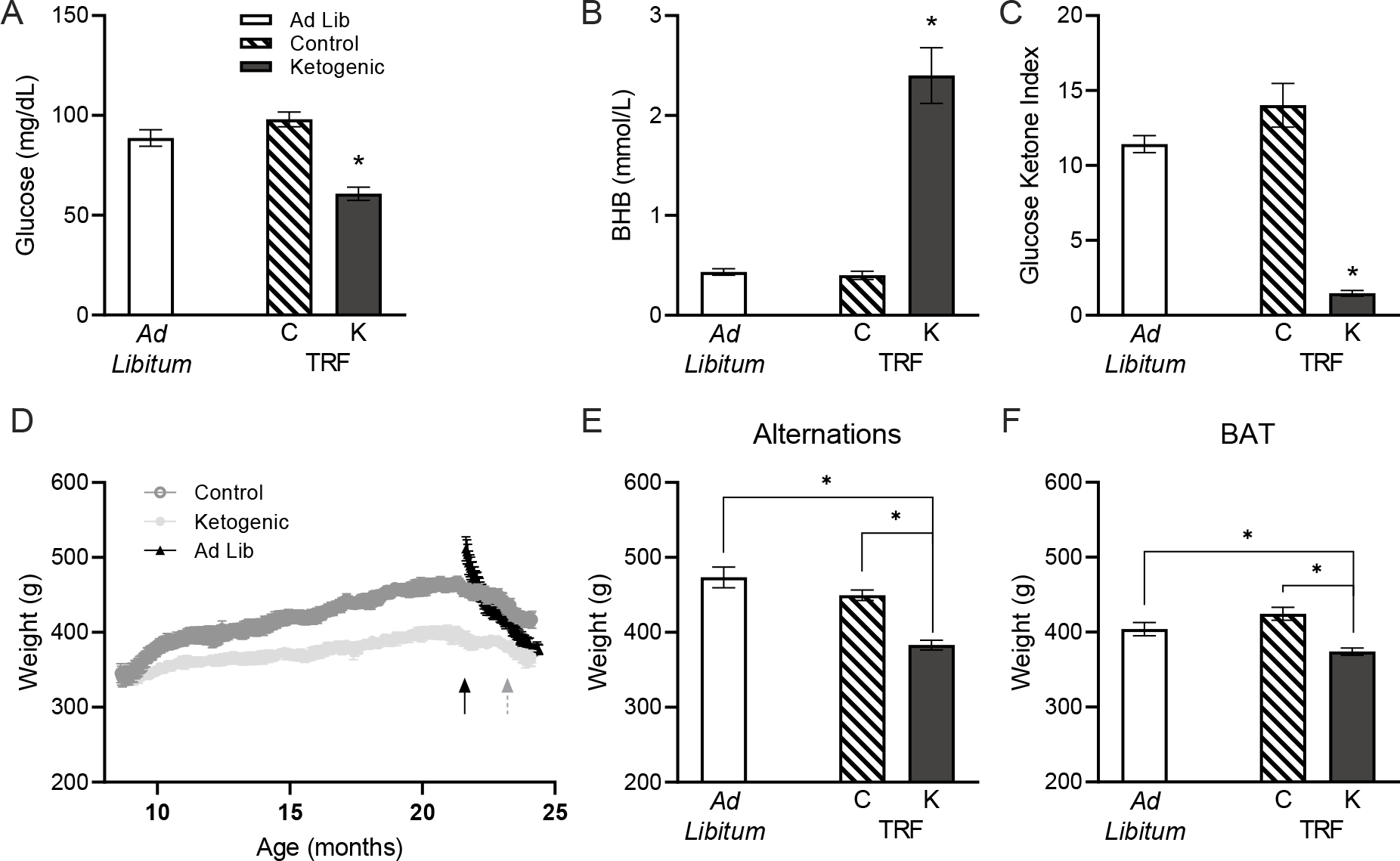
Postprandial glucose, BHB and GKI values and body weight. Only rats fed a ketogenic diet exhibited nutritional ketosis as evidenced by (A) reduced glucose, (B) elevated levels of the ketone body β-hydroxybutyrate and (C) a lower glucose ketone index (GKI). (D) Body weights throughout the lifespan continued to increase while fed a ketogenic or control diet via time-restricted feeding, until the onset of further dietary restriction (solid black arrow) and during BAT testing (beginning at dashed gray arrow). Body weight on the first day of (E) alternation training, as well as (F) BAT, was significantly lower in ketogenic diet-fed rats relative to both other groups. All data are group means ± 1 SEM; * indicates p < 0.05.

Over the course of time-restricted feeding, both ketogenic and control-fed rats gained a significant amount of weight (F_[1,6]_ = 267.50; p < 0.001), indicating that they were not calorically restricted. Although caloric intake was identical, the control-fed rats gained significantly more weight that the ketogenic diet-fed rats (F_[1,6]_ = 7.77; p = 0.03). Moreover, the interaction between time and diet group was significant (F_[1,6]_ = 44.30; p = 0.001), indicating that the control-fed rats also gained weight more rapidly. Although this observation suggests that rats were not calorically restricted, weights did not reach the level of *ad libitum*-fed animals of the same age (Figure 2D). Body weight on the first day of alternation and BAT testing was significantly lower in ketogenic diet-fed rats than control-fed (t_[6]_ = 6.86; p < 0.001; t_[6]_ = 5.04; p = 0.002 respectively) and *ad libitum*-fed rats (t_[5]_ = 6.42; p = 0.001; t_[5]_ = 3.24; p = 0.02 respectively), though the control-fed and *ad libitum*-fed rats did not significantly differ (t_[5]_ = 1.62; p = 0.17; t_[5]_ = 1.65; p = 0.16; respectively Figures 2D-E). The comparable body weights during behavioral testing between the time-restricted control-fed rats and rats that ate *ad libitum* between 8 and 21 months suggest that potential differences in behavior cannot be explained by differences in overall body condition.

The total number of incorrect trials across all days of training through the final day of criterion performance for all tasks were tabulated for each rat. For alternation training, neither diet group (F_[2,8]_ = 1.88; p = 0.21) nor feeding method (F_[1,9]_ = 4.23; p = 0.07) significantly affected the number of incorrect trials required to reach criterion performance on alternations throughout the maze, though there was a trend for rats that were on *ad libitum* feeding prior to testing to make more errors during alternation training (Figure 3A).

**Figure 3:**
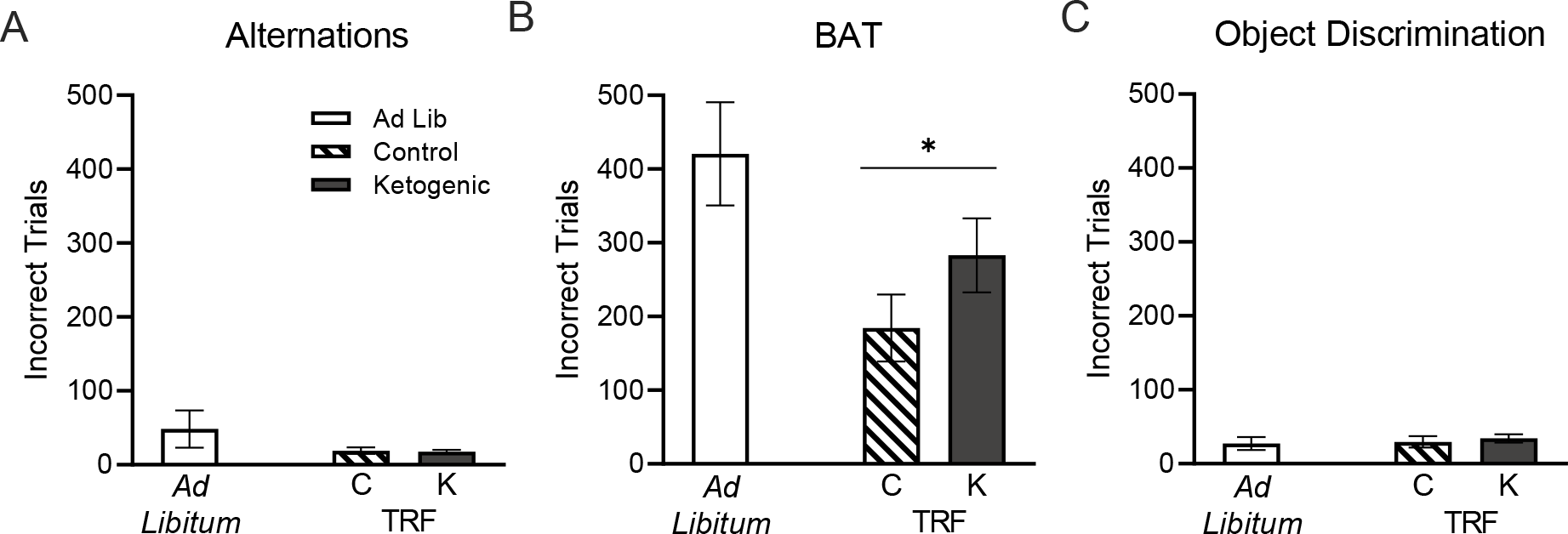
Behavioral performance of aged rats across diet groups and feeding methods. (A) There were no differences across groups in ability to alternate between the left and right arms of the maze. (B) Lifelong time-restricted feeding, regardless of macronutrient composition, improved ability to acquire the object-in-place rule required for criterion performance on the BAT task. (C) All rats were able to perform an object discrimination task similarly, indicating no physiological differences across groups that would prevent BAT task performance. All data are group means ± 1 SEM; * indicates p < 0.05.

The number of incorrect trials required to reach criterion performance on BAT testing was significantly different across the diet groups (F_[2,8]_ = 4.55; p = 0.048; Figure 3B). These results were due to the time-restricted feeding from young adulthood rather than dietary macronutrient composition, as the method of feeding also had a significant effect on performance (F_[1,9]_ = 6.62; p = 0.03). Conversely, among the rats that were given time-restricted feeding from young adulthood into old age, there was no difference in performance between the ketogenic and control diet groups (t_[6]_ = 1.46; p = 0.20).

Following BAT testing, a simple object discrimination control was utilized to ensure all rats were able to discriminate between two dissimilar objects and to assess potential differences in motivation or procedural impairments (Figure 3C). There was not a significant difference in the number of incorrect trials required to reach a criterion performance on this task across the 3 groups of rats (F_[2,8]_ = 0.23; p = 0.80), nor was there an effect of feeding method (*ad libitum* versus time-restricted; F_[1,9]_ = 0.25; p = 0.63). These data demonstrate all rats were able to discriminate between objects and are not visually impaired, indicating BAT task performance was not hindered by physical or sensory deficits, but rather by than cognitive differences across diet groups.

## Discussion

The data presented here show that time-restricted feeding initiated in mature adults, prior to the onset of age-related metabolic impairments, can influence cognitive outcomes in advanced age. Specifically, rats fed once daily between the ages of 8 and 21 months with either a ketogenic or a standard control diet performed better on a cognitive biconditional association task compared to rats that were fed *ad libitum* during that same time frame. This observation could not be accounted for by differences in body weight or sensorimotor impairments. It is important to note than although rats were fed once daily, and thus prevented from obesogenic overconsumption typically observed with *ad libitum* feeding (13), these rats were not calorically restricted and continued to show modest increases in weight throughout their lives.

Furthermore, we observed that nutritional ketosis, which has been shown to improve metabolic health in aged rats (5,6), did not confer an additive benefit to time-restricted feeding with a standard control diet in regards to multitasking performance on a biconditional association task. Shorter term ketogenic diets initiated in old age (21 months), which is an age at which rats on lifelong *ad libitum* feeding have age-related declines in metabolic function, have demonstrated improved cognitive outcomes on a similar task as well as reduced anxiety-like behavior (9). An important distinction between these two studies is the timing of the initiation of ketogenic diet therapy. When initiated at 8 months of age, normal rats show little metabolic dysfunction from lifelong *ad libitum* feeding with a standard diet. In contrast, rats that are allowed to consume food *ad libitum* into old age gain excessive weight, acquire aberrant amounts of white adipose tissue, as well as show disrupted insulin signaling and a reduced ability to utilize glucose in the brain (5,6,13,14). The current data suggest that time-restricted feeding throughout adulthood, regardless of the macronutrient composition, may be able to prevent these metabolic deficits in old age and lead to resilience against age-related cognitive decline. In contrast, when diet interventions are initiated in old age, declines in metabolic function need to be reversed. A previous study reported that time-restricted feeding with a ketogenic diet may be more effective at normalizing metabolic function in aged rats than time-restricted feeding with a standard diet. Thus, it may be critical to consider an individual’s current metabolic status when designing an optimal diet-based intervention for optimizing cognitive performance. This type of precision medicine-based approach has recently been suggested as an new avenue for treating cognitive aging (15).

The current data also have important implications in mid-life food consumption patterns and later life cognition. While is it well documented that high fat/high sugar obesogenic diets are associated with worse cognition (16), these data show that over consuming a standard diet that does not contain high fat or excessive sugar can also lead to worse to cognitive performance later in life. Thus, it is conceivable that adults who overconsume throughout mid-life are at a higher risk for cognitive decline in advanced age. A 2017 CDC study found that 44.7% of individuals aged 40-59 in the United States are obese (17), demonstrating the dire necessity of interventions during this critical period to avoid further cognitive decline in geriatric populations.

## Author contributions

ARH designed the study, contributed to data acquisition, analysis and interpretation, and prepared the manuscript. QPF, contributed to data acquisition and manuscript preparation. SNB designed the study, contributed to data analysis, interpretation and prepared the manuscript.

## Acknowledgements of Funding

This work was supported by the National Institutes of Health, National Institute on Aging (RF1AG060977; 1F31AG058455).

